# Shark: fishing in a sample to discard useless RNA-Seq reads

**DOI:** 10.1101/836130

**Authors:** Paola Bonizzoni, Tamara Ceccato, Gianluca Della Vedova, Luca Denti, Yuri Pirola, Marco Previtali, Raffaella Rizzi

## Abstract

Recent advances in high throughput RNA-Seq technologies allow to produce massive datasets. When a study focuses only on a handful of genes, most reads are not relevant and degrade the performance of the tools used to analyze the data. Removing such useless reads from the input dataset leads to improved efficiency without compromising the results of the study.

To this aim, in this paper we introduce a novel computational problem, called gene assignment and we propose an efficient alignment-free approach to solve it. Given a RNA-Seq sample and a panel of genes, a gene assignment consists in extracting from the sample the reads that most probably were sequenced from those genes. The problem becomes more complicated when the sample exhibits evidence of novel alternative splicing events.

We implemented our approach in a tool called `Shark` and assessed its effectiveness in speeding up differential splicing analysis pipelines. This evaluation shows that `Shark` is able to significantly improve the performance of RNA-Seq analysis tools without having any impact on the final results.

The tool is distributed as a stand-alone module and the software is freely available at https://github.com/AlgoLab/shark.

## 1. Introduction

RNA-Seq analysis plays an important role in the biological and medical research aimed at deepening our understanding of cellular biological processes and their relationships with pathological conditions. As such, several research initiatives had the objective of producing RNA-Seq data, while a number of tools have been proposed to analyze such datasets and gained a widespread adoption in the community [4, 8, 16, 18, 19, 24]. Traditional approaches generally rely on RNA-Seq read mapping to the annotated gene isoforms or on the spliced alignment of reads to the genome. In the second case, spliced aligners such as `STAR` [9] employ gene annotations to obtain more accurate results. While gene annotations, especially for humans, are readily available, they are not complete as they are built from healthy individuals. Thus, aberrant isoforms, which play an important role in the development of human diseases [12, 23], are usually not annotated. As a consequence, *de novo* (or assembly-first) approaches, which potentially detect novel splicing events, have been developed and gained popularity, even for studying well-annotated organisms. These approaches are more computationally demanding than traditional annotation-guided ones and pose challenging issues when facing the huge amount of RNA-Seq data that is now available. Furthermore, high-coverage samples are needed for obtaining accurate results with *de novo* approaches, especially for reconstructing low-abundance isoforms.

On the other hand, if the study is focused on the analysis of a pre-identified set of genes — for example, those that are known to have a role in tumor progression — then a preprocessing step that filters the input RNA-Seq reads, retaining only those likely originated by the genes of interest, could greatly reduce the size of the dataset that must be analyzed (hence speeding up the analysis) without significantly affecting the final results. Existing spliced aligners, such as `STAR`, could be theoretically adapted to perform the preprocessing step. However, they are aimed at obtaining accurate alignments, hence they are not fast enough to give a significant speedup of the analysis. Other approaches, such as some recently proposed transcript quantification tools [4, 16, 17] or quasi-mappers [20], are fast enough to be adapted as a preprocessing filters but they rely on isoform annotations, hence they might make errors if novel splicing events are supported by the sample, influencing any downstream analysis [5].

For this reason, we propose an alignment-free method that solves the gene-read assignment problem without relying on existing isoform annotations: given a set of genes of interest and a genome-wide RNA-Seq dataset, the goal is to retain only those reads (called *relevant* reads) originating from a gene in the selected set, thus discarding a potentially huge set of reads not relevant for the downstream analyses. We have implemented the method in the tool `Shark` that, by relying on succinct data structures and multi-threading, is able to quickly analyze huge RNA-Seq datasets on a standard PC. We performed an experimental evaluation on real data to assess the accuracy and the efficiency of our tool, showing that it achieves our goal of providing a preprocessing step that significantly reduces the running time of computationally intensive downstream analyses, while not negatively impacting their the results.

More precisely, we considered the following three pipelines: `STAR` followed by `SplAdder, STAR` followed by `rMATS`, and `Salmon` followed by `SUPPA2`. To perform this analysis, we considered a set of 6 RNA-Seq samples (three replicates for two conditions) produced by the study proposed in [2] where 83 alternative splicing events have been validated experimentally by RT-PCR. We ran the three aforementioned pipelines providing the input RNA-Seq samples as well as the RNA-Seq samples produced by filtering with `Shark` the six samples with respect to the genes involved in the RT-PCR validated events. The results show that `Shark` is able to significantly reduce the size of the input RNA-Seq sample without losing information that is fundamental in downstream analysis. Indeed, `Shark` allows to speed up the three pipelines without affecting negatively their accuracy in detecting the RT-PCR validated events differentially expressed between the two conditions.

## 2. Preliminaries

Let Σ be a finite alphabet of size *σ* and let *s* = *c*_1_,…, *c_n_* be an ordered sequence of *n* characters drawn from Σ; we say that *s* is a string over Σ of length n. From a computational point of view, a *genome*, a *gene locus*, and a *transcript* are strings over the alphabet {`A, C, G, T`}. A gene locus is a substring of the genome, whereas a transcript is a concatenation of pieces (exons) of a gene locus; a RNA-Seq sample is a set of strings (called *reads*) over the same alphabet. A RNA-Seq read is a substring of a transcript, and it is in a 1-to-1 correspondence with a gene locus, referred in the following as the *origin* of the read. Given a string *s* and a positive integer *k*, we say that a substring of *s* of length *k* is a *k*-mer. We denote by `KMER`(*s*) the multiset of all the *k*-mers of *s* (observe that the same *k*-mer might occur multiple times). As usual [1, 7, 14, 15], to account for the double stranded nature of the human genome, when we refer to a *k*-mer, we implicitly refer to its *canonical form*, that is the lexicographically smaller sequence between the *k*-mer and its reverse-complement. Moreover, to avoid *k*-mers being equal to their reverse-complement, we will only consider odd values of *k*. Given a read *s* = *c_i_* … *c_n_*, we refer to the pair (*c_i_*, *i*) with the term *base*. Note that the same character appearing at two different positions of *s* are two distinct bases. Moreover, we say that a base *b* of the read *s* is shared with the gene *g* if there exists a *k*-mer of *s* that includes b and is equal to a *k*-mer of *g*. We denote by `SHARED`(*s, g*) the set of the bases of *s* shared by *g*. In other words, `SHARED`(*s, g*) is the set of all the bases of *s* such that there exists a *k*-mer in the intersection between `KMER`(*g*) and `KMER`(*s*) that includes *s*.

To assign a read to its origin gene, we adapt the following criterion as a proxy. A gene *g* is the putative origin of a read *s* if and only if the ratio |`SHARED`(*s, g*)|/|*s*| is greater or equal than a given threshold *τ* and for no other gene *g′*, |`SHARED`(*s, g′*)| > |`SHARED`(*s, g*)|. The rationale behind this criterion is to have a measure of similarity between a read and a gene without aligning them, which takes into account that introns are spliced out from transcripts and therefore from a RNA-Seq read. Observe that by this definition, and due to genome repetitions, a read may have multiple origin genes. We denote as `ORIGIN`(*s*) the set of putative origin genes of a read *s*.

We now provide a formal definition of the problem we tackle in this paper, namely the *Gene Assignment Problem*.

**Problem 1** (Gene Assignment Problem). Let *S* be a set of RNA-Seq reads, let *G* = {*g*_1_,…, *g_p_*} be a set of genes of interest. The *gene assignment* of *S* with respect to *G* and parameters *k* and *τ* is a set *A* = {*S*_1_,…, *S*_|*G*|_} of |*G*| elements such that *S_i_* ⊆ *S* is the set of reads that originate from *g_i_*, that is for each *s* in *S_i_* the following conditions hold: (1) |`SHARED`(*s, g_i_*)|/|*s*| ≥ *τ*, (2) for no other gene *g_j_*, |`SHARED`(*s, g_j_*)| > |`SHARED`(*s, g_i_*)|, and the `SHARED`(·, ·) sets have been computed on *k*-mers.

Note that a gene assignment is not necessarily a partition of the input sample. Indeed, a read may have more than one origin gene and some read may have no origin gene. For ease of presentation, in the rest of the paper we will refer to this problem as the gene assignment of *S* with respect to *G*, without specifying *k* and *τ*. The algorithmic approach we propose to solve this problem uses two well known data structures that we will now briefly introduce.

A *bit vector* is a sequence of binary values that supports `rank` and `select` queries in constant time using additional sublinear space. Let *B* be a bit vector and let *i* be a positive integer, `rank`*_d_*(*B, i*) is the number of values equal to *d* in the portion *B*[1, *i*) of *B*, whereas `select`_*d*_(*B, i*) is the position of the *i*-th value of *B* set to *d*. Clearly, `rank`_*d*_(*B, i*) is not defined if *i* is greater than the size of *B* and `select`_*d*_(*B, i*) is not defined if fewer than *i* values of *B* are set to *d*.

A *Bloom filter* [3] is a probabilistic data structure used to answer membership queries. It consists of a bit vector of fixed size *m* and *z* hash functions, each one mapping an object to a position of the bit vector. Adding an object to a Bloom filter consists in setting to 1 the bits at the positions of the bit vector computed by the hash functions for that object. Testing if an object is in the set consists in checking whether all the bits at the positions computed by the hash functions for that object are set to 1. Due to hash collisions, an element may be reported as present even though it is not in the set, resulting in a false positive. Anyway, a low false positive rate can be achieved via a suitable choice of the bit vector size and the number of hash functions.

## 3. Methods

In this section we describe the algorithmic approach we propose to solve the computational problem introduced in the previous section. The algorithm for computing the gene assignment of an RNA-Seq sample *S* with respect to a set *G* of genes is composed of two steps: first, for each read *s* in the sample we compute the set `ORIGIN`(*s*) of its origin genes, then we derive the gene assignment of *S* from those sets by grouping together reads with the same origin gene.

An efficient solution to this problem essentially requires that we index the gene sequences. A simple procedure stores a dictionary that maps each *k*-mer appearing in at least one gene to the genes in which it occur, and then use it to map the *k*-mers of the reads to the genes, determining the origin genes read by read. Although effective, this approach would require an excessive amount of memory to store the dictionary if we need to track a significant amount of *k*-mers of the genes, especially since we have to store explicitly the *k*-mers sequences. For this reason, we designed a novel data structure that couples efficient access with small space usage, albeit introducing some false positives.

The data structure we propose to efficiently compute a gene assignment consists of a Bloom filter *BF*, a bit vector *P*, and a vector of integers *I*. We use the three components of the data structure as follows: the Bloom filter stores the set of *k*-mers of the genes in *G*, the vector of integers compactly stores the subset of genes in which each *k*-mer appears, and the bit vector tags the boundaries of the different subsets in the integer vector.

To build this data structure we designed a 3-step process, also presented in Algorithm 1. First, we associate to each gene in *G* an incremental ID and store the gene-ID mapping in a dictionary `GENEMAP`, which is then given as argument to Algorithm 1. Notice that we achieve an important reduction in memory usage by storing only the gene ID instead of its entire sequence.

Then, we scan the gene sequences and we store each *k*-mer in *BF* by using a single hash function *H*, mapping each *k*-mer to a specific position in the Bloom filter *BF* (lines 1–5). Once we complete this scan, *BF* stores the set of *k*-mers of all input genes.

For each 1 in *BF*, we create an empty list *L_r_* (lines 6–7), where *L_r_* corresponds to the r-th bit stored in the Bloom filter. We will then use *L_r_* to store a set of back-references to the genes where each associated *k*-mer appears. At the end of this step, each 1 in *BF* is associated with a subset of genes back-references (represented as a list of IDs) stored in memory. Then, we scan all the *k*-mers in each gene of *G*, we compute the corresponding 1 in *BF* and (via a rank) the corresponding list *L_r_* to which the gene ID must be appended (lines 8–12). Finally all duplicates are removed from the lists *L_r_*.

The third step consists of concatenating the lists *L_r_* to obtain the integer vector *I* that contains all back-references. At the same time (lines 15–19), we build the boundary vector *P* which has a 1 in each position 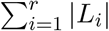 where *r* ≤ *ℓ* (i.e., the 1s mark the end of each list).

The example in Figure 1 describes how this data structure is queried to retrieve the identifiers of the genes associated to a given *k*-mer. First, the hash function *H* maps the 5-mer *gactgg* to position *h*. Since a 1 is at position *h* of *BF*, we suppose that the *k*-mer is in it and we compute how many 1s appear before *h* computing in constant time `rank`_1_(*BF, h*). Then, since the 1 stored in *h* is the *v*-th 1 of *BF*, *i.e*., the *k*-mer is the *v*-th element according to the order of *k*-mers given by *H*, we retrieve the positions of the (*v* − 1)-th and the *v*-th 1 in *P* using `select`_1_. Those positions are the boundaries on *I* of the subset of genes mapped to the *k*-mer.

**Figure 1:**
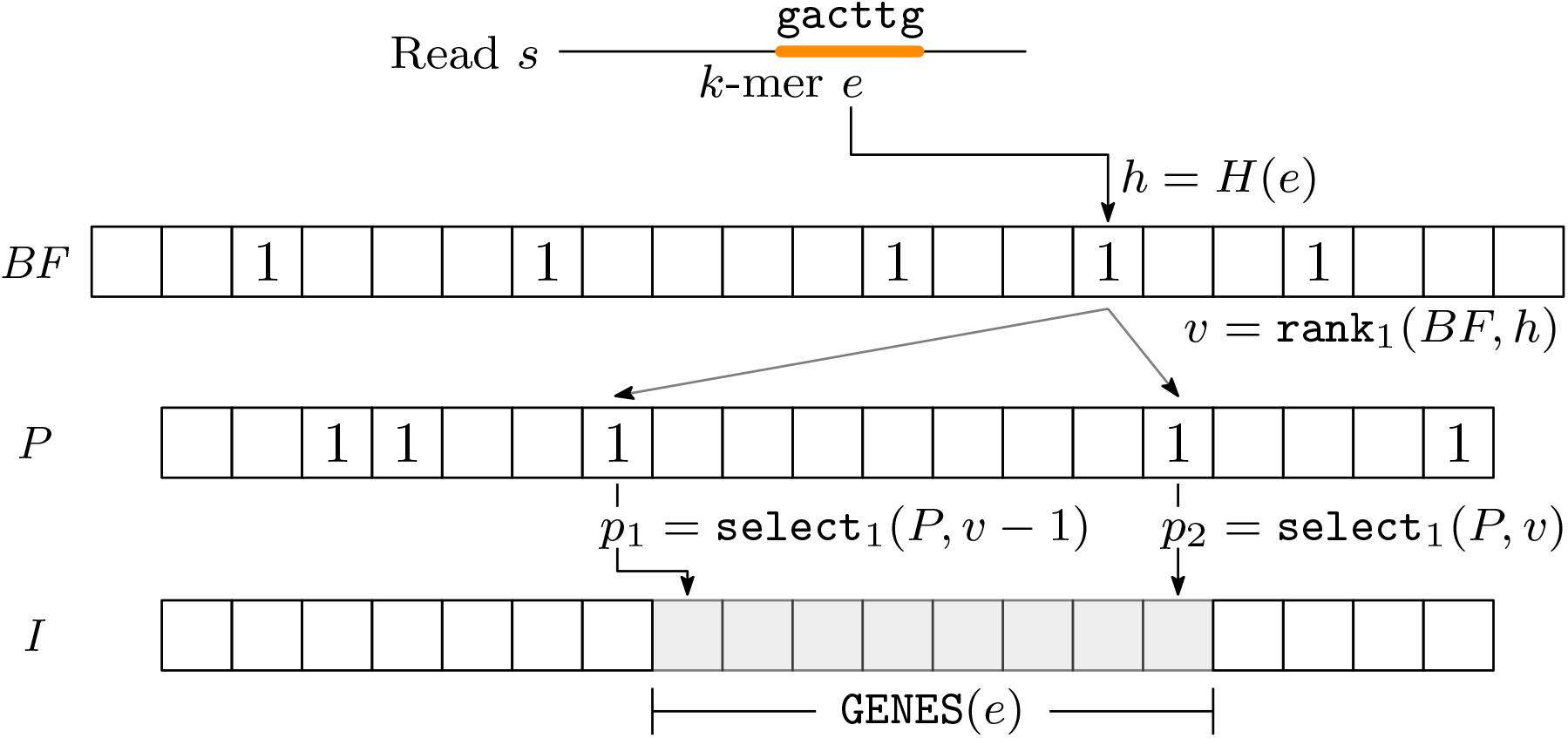
Relation between the Bloom filter *BF*, the bit vector *P*, and the vector *I*. To retrieve the identifiers of the genes containing a *k*-mer *e* (`gacttg` in the figure), we compute its image *h* through *H* and, if *BF*[*h*] is the *v*-th 1 of *BF*, the positions of the (*v* − 1)-th and the *v*-th 1 of *P*, denoted as *p*_1_ and *p*_2_ respectively, can be found via rank and select operations. The interval of *I* from *p*_1_ + 1 to *p*_2_ stores the set `GENES`(*e*) of the indices of the genes containing the *k*-mer *e*.

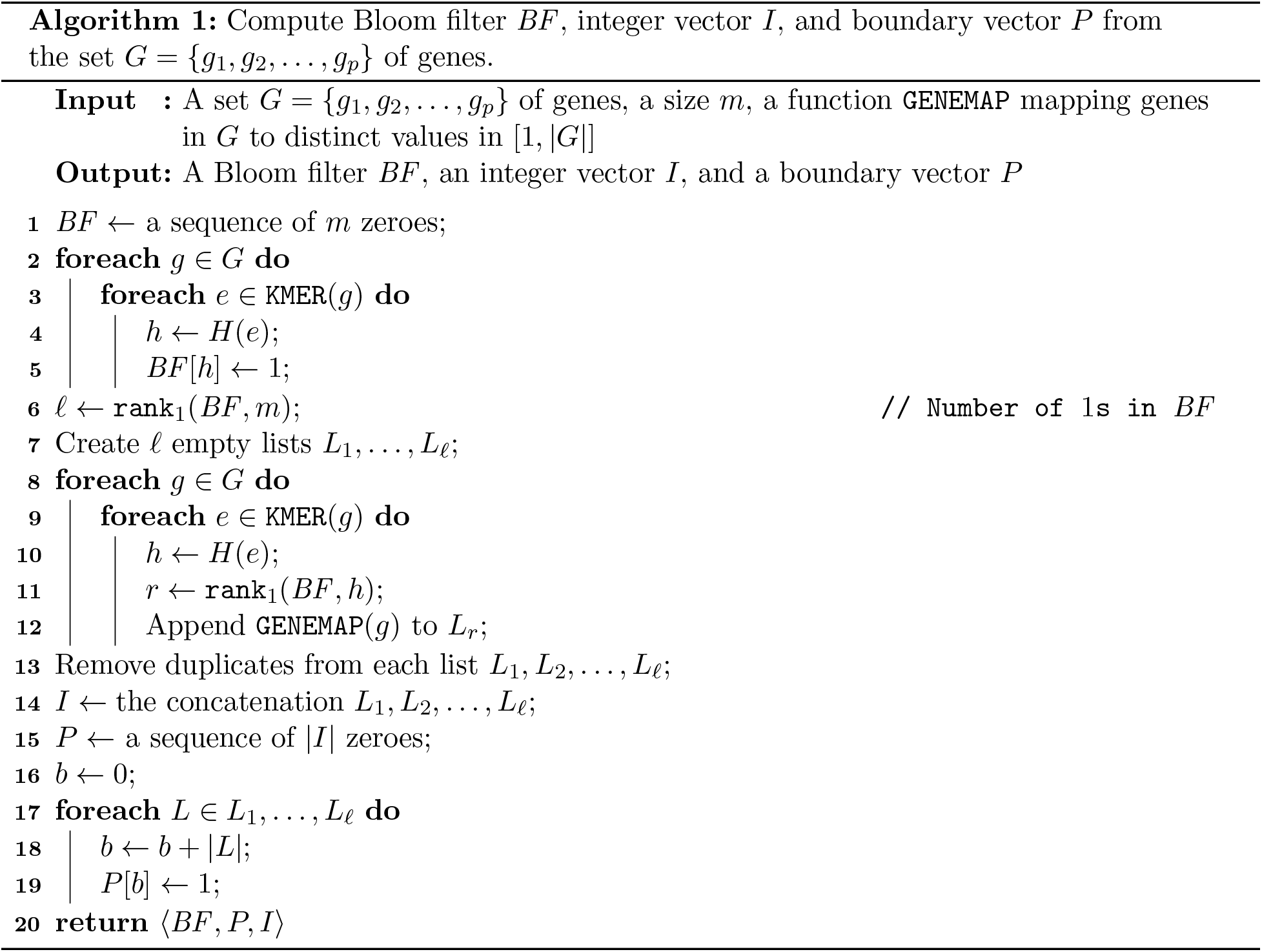

Once we have built the data structure *D*(*G*) = 〈*BF,P,I*〉, as described above, we iterate over each read of the sample and we query *D*(*G*) to compute the set of its origin genes. For each read *s*, the procedure scans the multiset `KMER`(*s*) of *k*-mers of *s* from left to right and maintains, for each gene *g*, the rightmost position *pos_g_* of a base of *s* shared by *g* (or −1 if such a base does not exist) and the variable *card_g_* that contains the number of bases of *g* that are shared with some *k*-mers of *s* seen so far — at the end, *card_g_* will be equal to |`SHARED`(*s, g*)|. Let e be the *i*-th leftmost *k*-mer of *s* and let `GENES`(*f*) be the set of genes containing *e* (obtained by querying *D*(*G*) as illustrated in Figure 1). Then, for each gene *g* in `GENES`(*e*), the procedure updates *card_g_* by adding the number of new bases covered by *e* and shared between *s* and *g* (computed as min{*k, i* + *k* − *pos_g_*}), and sets *pos_g_* to the last position in *s* that is in *e*, equal to *i* + *k*. At the end of the scan, each *card_g_* is clearly the cardinality of `SHARED`(*s, g*) and computing the origin gene of the read is trivial.

## 4. Experimental results

The method described in Section 3 has been implemented in `C++` and is freely available at https://github.com/AlgoLab/shark. Such a tool, called `Shark`, takes as input a `FASTA` file containing the set of gene regions of interest and an RNA-Seq sample in `FASTQ` format. For each read of the sample, the tool computes its set of (putative) origin genes (computing the gene assignment is then trivial). It is possible to tune the computation of the gene assignment by setting the following parameters: the *k*-mer size *k*, the confidence *τ*, and the size *m* of the Bloom filter. The tool also allows to filter out bases whose quality is less than a given threshold *q*.

The program uses the implementation of bit vectors and of the associated rank and select operations provided by `sdsl` [10]. As described in Section 3, the Bloom filter was implemented as the union of a bit vector and a single hash function. As proved by various works[7, 21, 22], in most cases using a single hash function has similar results as using multiple ones.

To assess our method, we performed two different experimental analyses on simulated and real data. A first exploratory analysis on simulated data was performed to test the accuracy and the efficiency of `Shark`, especially with respect to its input parameters *k*, *τ*, and *q*. The goal of the second analysis was to evaluate, on real data, the effectiveness of `Shark` in speeding up three different pipelines for a common analysis of RNA-Seq data, that is the differential analysis of alternative splicing events, without affecting the results on the selected genes. All the experiments were performed on a 64 bit Linux (Kernel 4.4.0) system equipped with four 8-core Intel^®^ Xeon 2.30GHz processors and 256GB of RAM.

### 4.1. Simulated data

We performed an exploratory analysis on simulated data to test the accuracy and the efficiency of our method, especially with respect to the input parameters *k* and *τ*. To this aim, we considered the 9403 genes of Human chromosomes 1, 17, and 21 *(Ensembl release 97* [6]) and we simulated an RNA-Seq sample of 10 million 100bp-long single-end reads using `Flux Simulator` [11]. From the full set of genes, we selected 10 random subsets of 100 genes, producing 10 different instances. To assess accuracy and efficiency of `Shark` with respect to input parameters, we ran `Shark` with any combination of *k* ∈ {13, 17, 23, 27, 31}, *τ* ∈ {0.2, 0.4, 0.6, 0.8}, and *q* ∈ {0, 10, 20}. Furthermore, we tested `Shark` also by dropping reads that are assigned to more than one origin gene (“single mode”). The size of the Bloom filter was set to 1GB (preliminary experiments showed that larger Bloom filters did not improve the accuracy of the prediction). `Shark` was allowed to use 4 threads to speed-up the computation.

The accuracy of `Shark` in computing the gene assignment was measured in terms of *precision*and *recall* as follows. Since the input reads have been simulated, we know the actual origin gene of each read. We considered each read associated to its origin gene as a *true positive* (*tp*), each read assigned to a gene different from the one it was simulated from as a *false positive* (*fp*), and each read simulated from one of the considered genes and not assigned to the right gene as a *false negative* (*fn*) Notice that when a read simulated from one of the 100 genes is assigned to the wrong gene, it induces both a false positive (because it was assigned to the wrong gene) and a false negative (because it was not assigned to the right gene). Then, precision is defined as 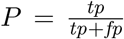, while recall as 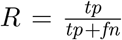. Efficiency was measured in terms of running time and memory peak (using the `/usr/bin/time` system tool).

Table 1 reports a representative subset of the results of this analysis. (See Table 4 in the Supplementary materials for the entire set of results.) The first important observation is that `Shark` achieves a good accuracy in terms of recall. Indeed, for sensible choices of the parameters, the recall is at least 99%, *i.e*., on average at most 1% of the reads that originated from the chosen subset of genes was discarded. The second observation is that the choice of the values for the parameters *k* and *τ* allows to achieve different trade-offs between precision and recall. Indeed, as *k* (or *τ*) increases, `Shark` becomes more precise (at the expense of recall).

The third observation is that `Shark` (as expected, since it is a *k*-mer-based approach) is sensitive to sequencing errors. Indeed, if low quality bases are not filtered out (*q* = 0), the recall rapidly decreases under 98% as *τ* increases. However, the simple approach of filtering out low quality bases allows to reduce the loss of recall as the precision sensibly increases. For example, for *q* = 10 (*i.e*., bases with quality less than 10 are not considered) and *k* = 17, when *τ* increases from 0.4 to 0.6, we have that the precision gains 7.13 percentage points (from 21.67% to 28.80%) while the recall only decreases of 0.21 percentage points (from 99.67% to 99.46%). Interestingly, an aggressive low-quality filter (*q* = 20) generally decreases the accuracy of `Shark`. This is probably due to the fact that, under this setting, the set of *k*-mers extracted from each read is too small for reliably finding its origin gene.

As explained in Section 3, more than one gene can be assigned to a single read. To assess how these ambiguous assignments influence the accuracy, we ran `Shark` excluding any multiple assignment (“single mode”). The accuracy of `Shark` in “single mode”compared with the original “multiple mode” is sensibly higher in terms of precision and slightly lower in terms of recall. For example, for *k* = 17, *τ* = 0.6, and *q* = 10, in multiple mode the precision is 28.8% while in single mode it is 32.2%, whereas the recall is 99.46% and 99.28% in multiple and single mode, respectively. Also in this case, keeping or discarding ambiguous assignments is an user’s choice, depending on the choice between higher recall or higher precision.

**Table 1:** Accuracy and efficiency results – Exploratory analysis. Accuracy is shown in terms of precision and recall whereas efficiency in terms of time (seconds). Precision, Recall, and Time are the average results obtained across the 10 performed runs.

| $k$ | $\tau$ | $q$ | Multiple mode | | | Single mode | | |
| --- | --- | --- | --- | --- | --- | --- | --- | --- |
|  |  |  | Precision | Recall | Time (s) | Precision | Recall | Time (s) |
| 17 | 0.2 | 0 | 13.72 | 99.68 | 38 | 16.82 | 99.52 | 40 |
|  |  | 10 | 12.29 | 99.74 | 38 | 15.54 | 99.54 | 37 |
|  |  | 20 | 11.87 | 99.65 | 34 | 16.51 | 99.45 | 35 |
|  | 0.4 | 0 | 22.62 | 99.19 | 39 | 25.28 | 99.04 | 39 |
|  |  | 10 | 21.67 | 99.67 | 38 | 25.09 | 99.49 | 35 |
|  |  | 20 | 21.61 | 99.47 | 34 | 26.72 | 99.28 | 35 |
|  | 0.6 | 0 | 30.04 | 97.71 | 39 | 32.49 | 97.54 | 40 |
|  |  | 10 | 28.80 | 99.46 | 36 | 32.20 | 99.28 | 36 |
|  |  | 20 | 29.80 | 98.37 | 33 | 34.77 | 98.20 | 34 |
| 23 | 0.2 | 0 | 12.71 | 99.56 | 36 | 16.68 | 99.42 | 36 |
|  |  | 10 | 12.76 | 99.53 | 34 | 17.22 | 99.35 | 34 |
|  |  | 20 | 12.74 | 99.16 | 32 | 19.34 | 98.98 | 33 |
|  | 0.4 | 0 | 28.04 | 98.93 | 36 | 32.58 | 98.79 | 36 |
|  |  | 10 | 26.74 | 99.43 | 34 | 32.48 | 99.28 | 33 |
|  |  | 20 | 27.29 | 98.34 | 31 | 35.51 | 98.17 | 31 |
|  | 0.6 | 0 | 38.25 | 97.09 | 34 | 41.66 | 96.93 | 34 |
|  |  | 10 | 37.02 | 98.81 | 34 | 41.54 | 98.66 | 32 |
|  |  | 20 | 39.29 | 94.37 | 30 | 45.57 | 94.17 | 31 |
| 27 | 0.2 | 0 | 14.68 | 99.37 | 36 | 19.14 | 99.22 | 34 |
|  |  | 10 | 14.87 | 99.29 | 32 | 19.88 | 99.14 | 33 |
|  |  | 20 | 15.30 | 98.42 | 30 | 22.81 | 98.24 | 30 |
|  | 0.4 | 0 | 32.16 | 98.66 | 33 | 37.80 | 98.52 | 33 |
|  |  | 10 | 30.25 | 99.15 | 32 | 37.24 | 98.99 | 33 |
|  |  | 20 | 31.44 | 96.81 | 30 | 41.01 | 96.65 | 30 |
|  | 0.6 | 0 | 44.00 | 96.29 | 33 | 48.15 | 96.15 | 33 |
|  |  | 10 | 42.96 | 97.88 | 32 | 48.13 | 97.71 | 32 |
|  |  | 20 | 45.60 | 90.00 | 29 | 52.17 | 89.85 | 29 |
| 31 | 0.2 | 0 | 16.73 | 99.06 | 33 | 21.73 | 98.91 | 34 |
|  |  | 10 | 17.10 | 98.94 | 30 | 22.72 | 98.77 | 30 |
|  |  | 20 | 18.46 | 96.96 | 29 | 26.46 | 96.78 | 29 |
|  | 0.4 | 0 | 35.63 | 98.33 | 34 | 42.58 | 98.16 | 35 |
|  |  | 10 | 33.28 | 98.70 | 32 | 41.51 | 98.57 | 32 |
|  |  | 20 | 34.84 | 94.82 | 29 | 45.48 | 94.66 | 30 |
|  | 0.6 | 0 | 51.68 | 95.18 | 34 | 55.79 | 95.05 | 32 |
|  |  | 10 | 50.50 | 96.49 | 31 | 55.71 | 96.37 | 32 |
|  |  | 20 | 53.37 | 84.60 | 28 | 59.46 | 84.44 | 29 |

In terms of computational requirements, `Shark` never required more than 2.1GB of RAM, that is an amount of memory nowadays available on any desktop or laptop computer, and it never required more than 2 minutes to complete (using only 4 threads). Albeit this experiment has been performed on a server platform, we expect that the running time will be practically negligible even on standard computers.

This experiment showed how accuracy is influenced by the choice of the parameters. The results suggest that a good trade-off between precision and recall can be achieved with *k* = 17, *τ* ∈ {0.4,0.6}, and *q* = 10.

**Table 2:**
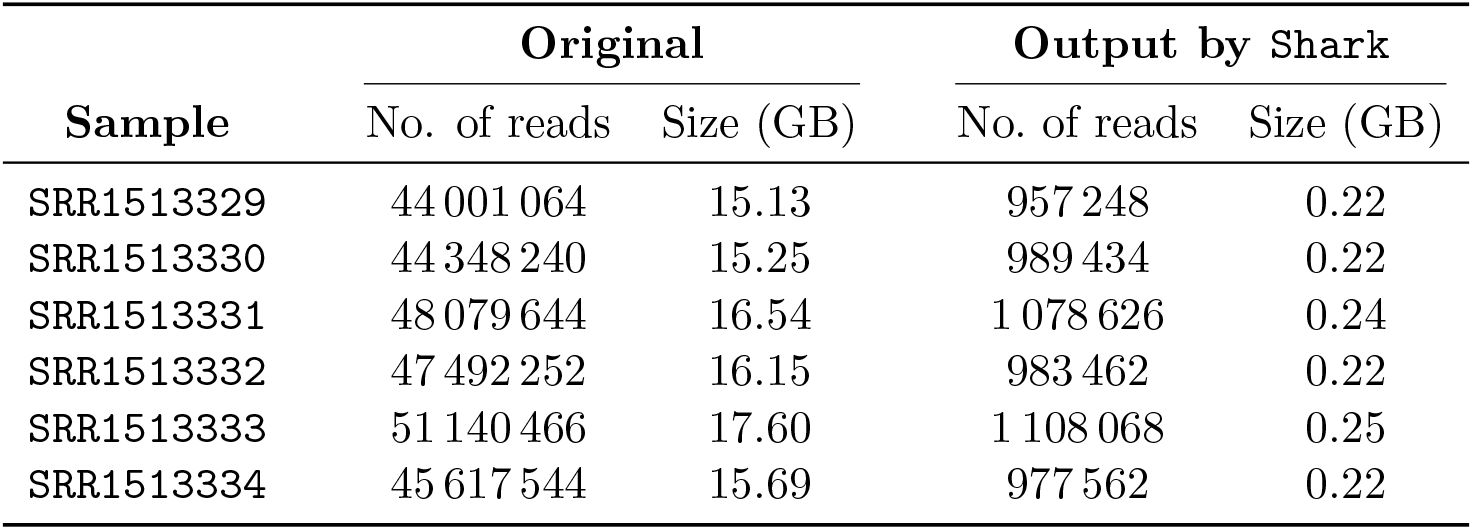
Sizes of the RNA-Seq samples considered in our experimental analysis on real data. The table shows the sizes in terms of number of reads and the uncompressed file size of the 6 original samples and the 6 samples produced by `Shark` after filtering them with respect to the 82 genes involved in the 83 RT-PCR validated events.

| Sample | Original |  | Output by Shark |  |
| --- | --- | --- | --- | --- |
|  | No. of reads | Size (GB) | No. of reads | Size (GB) |
| SRR1513329 | 44 001 064 | 15.13 | 957 248 | 0.22 |
| SRR1513330 | 44 348 240 | 15.25 | 989 434 | 0.22 |
| SRR1513331 | 48 079 644 | 16.54 | 1 078 626 | 0.24 |
| SRR1513332 | 47 492 252 | 16.15 | 983 462 | 0.22 |
| SRR1513333 | 51 140 466 | 17.60 | 1 108 068 | 0.25 |
| SRR1513334 | 45 617 544 | 15.69 | 977 562 | 0.22 |

### 4.2. Real dataset

In the second part of our experimental evaluation we partially replicate the analysis of a real RNA-Seq dataset performed in [24] in order to assess the effectiveness of our tool in speeding up state-of-the-art pipelines for differential analysis of alternative splicing.

We considered the three pipelines based on `SplAdder` [13], `rMATS` [19], and `SUPPA2` [24]. The first two tools analyze the RNA-Seq alignments computed by a spliced aligner, while the last one –`SUPPA2`– analyzes the transcript quantifications computed by `Salmon`. In the following, we will refer to the three pipelines only by the name of the tools for the differential analysis, *i.e*., `SplAdder`, `rMATS`, and `SUPPA2`. We remark that the aim of this part is not to evaluate the accuracy of the results of these pipelines, but to verify (1) whether their findings are affected by the preprocessing step performed by `Shark` and (2) how much `Shark` can speedup their analyses.

The dataset used in this evaluation consists of a set of six paired-end RNA-Seq samples (GEO accession number *GSE59335*): three samples obtained before and three obtained after the double knockdown of two splicing regulatory proteins, namely `TRA2A` and `TRA2B` [2]. Each sample contains between 44 and 51 millions of reads of length 101bp. We decided to test the tools on this dataset since in the study 83 exon skipping events have been validated experimentally by RT-PCR, thus we can use such events as a ground truth to assess the effect of using `Shark` as preprocessing step.

In this analysis we ran `Shark` setting *k* equal to 17, *τ* equal to 0.6, *q* equal to 10, and the Bloom filter size to 1GB. We chose these values since, as proved in the previous analysis, they achieve a good trade-off between precision and recall. All tools were ran with their default parameters (allowing up to 4 threads), while the spliced alignments required by the first two pipelines were computed by `STAR` [9] (in two-pass mode).

We initially computed the differentially spliced events with the three aforementioned pipelines considering the original RNA-Seq samples and then we repeated the analysis considering the RNA-Seq samples preprocessed with `Shark` on the 82 different gene regions involved in the 83 RT-PCR validated events. Table 2 reports the differences in terms of number of reads and uncompressed file size between the original samples and the samples filtered by `Shark`. On average, the filtered samples contain approximately the 2.3% of the original reads.

We considered the 83 alternative splicing events validated by RT-PCR and we evaluated if the ability of three pipelines in detecting such events is affected by the preprocessing step performed by `Shark`. Table 3 reports the results of this analysis.

The first observation is that all the pipelines detected the same RT-PCR validated events in both the considered scenarios *(i.e*., on the original samples and on the filtered ones), confirming that the preprocessing step performed with `Shark` does not affect the accuracy of their differential splicing analysis. More precisely, out of the 83 RT-PCR validated events, `rMATS` detected 78 differentially spliced events, `SUPPA2` detected 66 events, and `SplAdder` detected 56 events, under both the two conditions.

If we restrict our analysis only to events reported as statistically significant by each tool (*i.e*., the events with *p*-value smaller than 0.05), the results follow the same trend, further confirming that the preprocessing step does not negatively impact the differential analysis. Indeed, `rMATS` reported the highest number of events (63) followed by `SUPPA2`. Interestingly, `SUPPA2` identified 6 additional significant events when considering the samples preprocessed by `Shark` w.r.t. the 37 events detected on the original samples. A manual inspection of the events in the two scenarios revealed that the differences between the respective *p*-values were rather small. On the other hand, in both scenarios `SplAdder` reported all the events with a *p*-value close to 1 (hence not significant).

The second important observation is that, as expected, preprocessing the input samples with `Shark` makes the three pipelines faster. Indeed, `Shark` (which required less than 5 minutes to process each sample) allows all those pipelines to complete their analysis in around half the time. More precisely, `rMATS` took only 3 hours (saving more than 2 hours), `SplAdder` completed its analysis in 5 hours (saving more than 8 hours), and `SUPPA2` took only half an hour instead of an hour. The difference of the running times in the two scenarios is important (especially for `rMATS` and `SplAdder`), even on the relatively small dataset we considered here. Notice that the running time of `Shark` is linear in the number of input reads, and we only read once the set of reads. This fact implies that larger datasets (for example, with more replicates or across several conditions or with higher-coverage samples) should show an even larger (absolute) reduction of running times of the complete analyses. While `STAR` and `Salmon` perform an indexing procedure only once, `Shark` indexes the input gene regions for each sample — hence a more refined implementation that builds the index of the gene sequences only once could save even more time, especially for a larger number of samples. The details of the running times of the various steps of each pipeline are reported in Supplementary Table 5.

Lastly, this experiment shows that peak memory usage is almost unaffected by `Shark`. Indeed `Shark` required less than 4.4GB of RAM to process the input samples, which is significantly less than the peak memory usage of `rMATS` and `SplAdder` (33.9GB) and comparable to that of `SUPPA2` (4.3GB). In particular, the peak memory usage for `rMATS` and `SplAdder` is reached in the alignment step. In this step, `STAR` loads the entire genome index in memory, hence its memory usage is largely independent on the input sample size.

Overall, `Shark` proved to be a preprocessing step that do not affect negatively any downstream pipeline for the differential analysis of alternative splicing. On the other hand, it allows to significantly reduce the size of the input samples, hence speeding up the pipelines, especially those based on read alignment, without losing any valuable information essential for an accurate and sensitive downstream analysis.

## 5. Conclusions

In this work, we introduced the novel computational problem of computing the gene assignment of an RNA-Seq sample with respect to a set of genes. We also proposed an algorithmic approach to solve this problem and we implemented it into a tool, `Shark`. To the best of our knowledge, `Shark` is the first tool specifically designed for computing a gene assignment.

**Table 3:** Accuracy and efficiency of the three pipelines for differential analysis of alternative splicing on the original samples compared with those obtained on the samples filtered by `Shark`. Accuracy is evaluated in terms of the number of RT-PCR validated events detected by each pipelines (over a total of 83 RT-PCR validated events). Efficiency is evaluated in terms of running time and maximum memory usage.

| Pipeline | RT-PCR Events |  | Time | RAM |
| --- | --- | --- | --- | --- |
| | All | $p\text{-value} < 0.05$ | (min) | (GB) |
| rMATS | 78 | 63 | 308 | 33.9 |
| Shark + rMATS | 78 | 63 | 142 | 33.9 |
| SplAdder | 56 | - | 796 | 33.9 |
| Shark + SplAdder | 56 | - | 295 | 33.9 |
| SUPPA2 | 66 | 37 | 65 | 4.3 |
| Shark + SUPPA2 | 66 | 43 | 34 | 4.4 |

We performed an experimental analysis on real data where we evaluated the effectiveness of `Shark` in speeding up state-of-the-art pipelines for the differential analysis of alternative splicing. The results of our evaluation prove that `Shark` allows to speed up such pipelines without negatively affecting their accuracy and the final result they obtain. Moreover, `Shark` can be easily used on a standard desktop computer. However, the accuracy and the efficiency of `Shark` heavily depend on its parameters *k*, *τ*, and *q* that are the *k*-mer size, the minimum confidence, and the base quality threshold, respectively. For this reason, future steps will focus on allowing `Shark` to automatically estimate the best values of these parameters by exploiting extra information on the read length and the error rate.

Since `Shark` can be used as a preliminary step in pipelines for the detection of novel alternative splicing events from samples of RNA-Seq data, future work will be devoted to an in-depth experimental analysis of `Shark` as a preliminary step of a pipeline that includes computationally-demanding tools such as `ASGAL` [8], that relies on mapping reads against a splicing graph, and `KISSPLICE` [18], that assembles reads and identifies alternative splicing events by analyzing bubbles in the resulting de Bruijn graph.

## A. Supplementary material

**Suppl. Table 4:** Accuracy and efficiency results – Exploratory analysis. Accuracy is shown in terms of precision and recall whereas efficiency in terms of time (seconds). Precision, Recall, and Time are the average results obtained across the 10 performed runs. The standard deviation is shown in brackets nearby each value.

| $k$ | $\tau$ | $q$ | Multiple mode | | | | | | Single mode | | | | | |
| --- | --- | --- | --- | --- | --- | --- | --- | --- | --- | --- | --- | --- | --- | --- |
| | | | Precision ($\sigma$) | | Recall ($\sigma$) | | Time ($\sigma$) | | Precision ($\sigma$) | | Recall ($\sigma$) | | Time ($\sigma$) | |
| 13 | 0.2 | 0 | 1.47 | (0.67) | 99.46 | (0.22) | 67 | (8.40) | 1.61 | (0.78) | 99.29 | (0.36) | 65 | (6.50) |
|  |  | 10 | 1.43 | (0.67) | 99.72 | (0.08) | 67 | (7.40) | 1.61 | (0.78) | 99.50 | (0.28) | 63 | (5.80) |
|  |  | 20 | 1.51 | (0.70) | 99.72 | (0.08) | 60 | (6.00) | 1.75 | (0.81) | 99.48 | (0.30) | 58 | (5.40) |
|  | 0.4 | 0 | 3.53 | (1.73) | 99.08 | (0.21) | 65 | (9.20) | 3.61 | (1.81) | 98.91 | (0.35) | 62 | (4.70) |
|  |  | 10 | 3.42 | (1.66) | 99.71 | (0.09) | 62 | (8.60) | 3.54 | (1.75) | 99.47 | (0.32) | 61 | (8.90) |
|  |  | 20 | 3.97 | (1.94) | 99.66 | (0.05) | 56 | (6.90) | 4.18 | (2.07) | 99.42 | (0.34) | 54 | (6.90) |
|  | 0.6 | 0 | 11.23 | (5.86) | 97.83 | (0.21) | 62 | (9.40) | 11.62 | (6.21) | 97.65 | (0.35) | 61 | (7.40) |
|  |  | 10 | 10.74 | (5.50) | 99.56 | (0.15) | 57 | (4.00) | 11.31 | (5.95) | 99.35 | (0.35) | 55 | (9.70) |
|  |  | 20 | 12.50 | (5.93) | 99.31 | (0.15) | 53 | (5.60) | 13.50 | (6.62) | 99.07 | (0.33) | 55 | (5.10) |
|  | 0.8 | 0 | 24.93 | (8.19) | 91.77 | (0.32) | 60 | (7.10) | 26.45 | (8.87) | 91.60 | (0.41) | 58 | (4.70) |
|  |  | 10 | 24.25 | (7.89) | 98.92 | (0.27) | 53 | (9.20) | 26.46 | (8.81) | 98.73 | (0.41) | 57 | (6.80) |
|  |  | 20 | 26.80 | (7.81) | 95.75 | (0.30) | 53 | (4.40) | 30.08 | (8.87) | 95.54 | (0.43) | 54 | (5.10) |
| 17 | 0.2 | 0 | 13.72 | (5.10) | 99.68 | (0.06) | 38 | (1.90) | 16.82 | (6.09) | 99.52 | (0.31) | 40 | (3.60) |
|  |  | 10 | 12.29 | (4.60) | 99.74 | (0.05) | 38 | (4.30) | 15.54 | (5.66) | 99.54 | (0.31) | 37 | (3.10) |
|  |  | 20 | 11.87 | (4.42) | 99.65 | (0.05) | 34 | (3.10) | 16.51 | (5.89) | 99.45 | (0.33) | 35 | (3.80) |
|  | 0.4 | 0 | 22.62 | (7.08) | 99.19 | (0.09) | 39 | (3.20) | 25.28 | (7.86) | 99.04 | (0.33) | 39 | (3.50) |
|  |  | 10 | 21.67 | (6.88) | 99.67 | (0.07) | 38 | (3.10) | 25.09 | (7.83) | 99.49 | (0.32) | 35 | (4.40) |
|  |  | 20 | 21.61 | (6.77) | 99.47 | (0.12) | 34 | (2.90) | 26.72 | (8.09) | 99.28 | (0.33) | 35 | (2.70) |
|  | 0.6 | 0 | 30.04 | (7.92) | 97.71 | (0.14) | 39 | (4.10) | 32.49 | (8.63) | 97.54 | (0.36) | 40 | (3.20) |
|  |  | 10 | 28.80 | (7.74) | 99.46 | (0.13) | 36 | (3.80) | 32.20 | (8.62) | 99.28 | (0.36) | 36 | (3.40) |
|  |  | 20 | 29.80 | (7.72) | 98.37 | (0.16) | 33 | (2.60) | 34.77 | (8.77) | 98.20 | (0.36) | 34 | (3.00) |
|  | 0.8 | 0 | 40.21 | (8.25) | 91.09 | (0.29) | 38 | (2.80) | 42.92 | (8.89) | 90.94 | (0.44) | 38 | (4.00) |
|  |  | 10 | 39.16 | (8.15) | 97.91 | (0.30) | 36 | (2.70) | 43.07 | (8.89) | 97.74 | (0.44) | 35 | (4.50) |
|  |  | 20 | 41.84 | (8.15) | 90.22 | (0.42) | 34 | (2.40) | 47.06 | (8.90) | 90.04 | (0.57) | 34 | (2.00) |
| 23 | 0.2 | 0 | 12.71 | (4.57) | 99.56 | (0.05) | 36 | (3.80) | 16.68 | (5.45) | 99.42 | (0.31) | 36 | (3.20) |
|  |  | 10 | 12.76 | (4.57) | 99.53 | (0.07) | 34 | (3.20) | 17.22 | (5.58) | 99.35 | (0.33) | 34 | (3.60) |
|  |  | 20 | 12.74 | (4.57) | 99.16 | (0.07) | 32 | (3.10) | 19.34 | (6.09) | 98.98 | (0.34) | 33 | (2.80) |
|  | 0.4 | 0 | 28.04 | (7.62) | 98.93 | (0.08) | 36 | (1.80) | 32.58 | (8.54) | 98.79 | (0.35) | 36 | (1.90) |
|  |  | 10 | 26.74 | (7.45) | 99.43 | (0.08) | 34 | (2.50) | 32.48 | (8.54) | 99.28 | (0.35) | 33 | (1.80) |
|  |  | 20 | 27.29 | (7.36) | 98.34 | (0.13) | 31 | (1.80) | 35.51 | (8.75) | 98.17 | (0.38) | 31 | (2.90) |
|  | 0.6 | 0 | 38.25 | (8.20) | 97.09 | (0.16) | 34 | (3.10) | 41.66 | (8.80) | 96.93 | (0.39) | 34 | (3.90) |
|  |  | 10 | 37.02 | (8.10) | 98.81 | (0.17) | 34 | (4.20) | 41.54 | (8.83) | 98.66 | (0.38) | 32 | (1.30) |
|  |  | 20 | 39.29 | (8.13) | 94.37 | (0.32) | 30 | (2.30) | 45.57 | (8.94) | 94.17 | (0.50) | 31 | (2.30) |
|  | 0.8 | 0 | 52.93 | (8.50) | 88.85 | (0.48) | 33 | (4.30) | 55.66 | (8.96) | 88.74 | (0.63) | 35 | (2.40) |
|  |  | 10 | 51.80 | (8.45) | 93.91 | (0.50) | 33 | (3.20) | 55.64 | (8.99) | 93.76 | (0.68) | 34 | (3.10) |
|  |  | 20 | 54.34 | (8.54) | 77.84 | (0.76) | 31 | (2.80) | 59.15 | (9.02) | 77.74 | (0.90) | 30 | (3.30) |
| 27 | 0.2 | 0 | 14.68 | (4.87) | 99.37 | (0.05) | 36 | (2.80) | 19.14 | (5.86) | 99.22 | (0.36) | 34 | (3.10) |
|  |  | 10 | 14.87 | (4.92) | 99.29 | (0.07) | 32 | (3.10) | 19.88 | (6.04) | 99.14 | (0.32) | 33 | (2.30) |
|  |  | 20 | 15.30 | (5.02) | 98.42 | (0.11) | 30 | (3.30) | 22.81 | (6.62) | 98.24 | (0.38) | 30 | (1.40) |
|  | 0.4 | 0 | 32.16 | (7.80) | 98.66 | (0.11) | 33 | (2.80) | 37.80 | (8.75) | 98.52 | (0.38) | 33 | (3.00) |
|  |  | 10 | 30.25 | (7.62) | 99.15 | (0.10) | 32 | (3.20) | 37.24 | (8.71) | 98.99 | (0.37) | 33 | (3.00) |
|  |  | 20 | 31.44 | (7.57) | 96.81 | (0.20) | 30 | (3.10) | 41.01 | (8.90) | 96.65 | (0.45) | 30 | (3.50) |
|  | 0.6 | 0 | 44.00 | (8.30) | 96.29 | (0.21) | 33 | (3.20) | 48.15 | (8.95) | 96.15 | (0.43) | 33 | (3.80) |
|  |  | 10 | 42.96 | (8.24) | 97.88 | (0.21) | 32 | (2.80) | 48.13 | (8.96) | 97.71 | (0.42) | 32 | (2.30) |
|  |  | 20 | 45.60 | (8.33) | 90.00 | (0.51) | 29 | (2.50) | 52.17 | (9.03) | 89.85 | (0.70) | 29 | (2.00) |
|  | 0.8 | 0 | 60.32 | (8.81) | 85.80 | (0.85) | 33 | (3.20) | 62.90 | (9.19) | 85.70 | (0.99) | 33 | (2.80) |
|  |  | 10 | 59.20 | (8.81) | 89.68 | (0.90) | 33 | (2.80) | 62.73 | (9.20) | 89.56 | (1.05) | 33 | (3.70) |
|  |  | 20 | 61.22 | (8.88) | 69.67 | (1.03) | 30 | (3.70) | 65.37 | (9.28) | 69.59 | (1.13) | 30 | (3.20) |
| 31 | 0.2 | 0 | 16.73 | (5.31) | 99.06 | (0.08) | 33 | (2.60) | 21.73 | (6.37) | 98.91 | (0.35) | 34 | (3.30) |
|  |  | 10 | 17.10 | (5.41) | 98.94 | (0.07) | 30 | (1.80) | 22.72 | (6.55) | 98.77 | (0.36) | 30 | (2.30) |
|  |  | 20 | 18.46 | (5.64) | 96.96 | (0.16) | 29 | (2.00) | 26.46 | (7.17) | 96.78 | (0.43) | 29 | (1.20) |
|  | 0.4 | 0 | 35.63 | (7.79) | 98.33 | (0.14) | 34 | (3.30) | 42.58 | (8.86) | 98.16 | (0.37) | 35 | (2.70) |
|  |  | 10 | 33.28 | (7.61) | 98.70 | (0.13) | 32 | (3.20) | 41.51 | (8.79) | 98.57 | (0.37) | 32 | (1.90) |
|  |  | 20 | 34.84 | (7.69) | 94.82 | (0.31) | 29 | (3.00) | 45.48 | (8.98) | 94.66 | (0.52) | 30 | (2.10) |
|  | 0.6 | 0 | 51.68 | (8.26) | 95.18 | (0.31) | 34 | (2.70) | 55.79 | (9.08) | 95.05 | (0.52) | 32 | (3.00) |
|  |  | 10 | 50.50 | (8.26) | 96.49 | (0.30) | 31 | (2.50) | 55.71 | (9.13) | 96.37 | (0.50) | 32 | (2.80) |
|  |  | 20 | 53.37 | (8.43) | 84.60 | (0.78) | 28 | (2.30) | 59.46 | (9.21) | 84.44 | (0.94) | 29 | (2.70) |
|  | 0.8 | 0 | 66.25 | (9.53) | 82.61 | (1.19) | 33 | (4.50) | 68.51 | (9.87) | 82.51 | (1.31) | 33 | (3.40) |
|  |  | 10 | 65.19 | (9.42) | 85.54 | (1.28) | 32 | (2.80) | 68.23 | (9.82) | 85.45 | (1.39) | 31 | (3.20) |
|  |  | 20 | 66.42 | (9.40) | 62.56 | (1.28) | 29 | (1.80) | 69.91 | (9.80) | 62.50 | (1.34) | 28 | (3.10) |

**Suppl. Table 5:** Running times (in seconds) of each step of the considered pipelines. The last columns show the gain in terms of seconds and hours of using `Shark` as preprocessing step over not using it.

| Pipeline | Sample | Shark | STAR |  | Salmon |  | Diff.<br>analysis | Total<br>time | Gain<br>(secs) | Gain<br>(h) | Gain<br>(%) |
| --- | --- | --- | --- | --- | --- | --- | --- | --- | --- | --- | --- |
| rMATS | SRR1513329 |  |  | 1 773 |  |  |  |  |  |  |  |
|  | SRR1513330 |  |  | 1 704 |  |  |  |  |  |  |  |
|  | SRR1513331 |  | 5 227 | 2 348 |  |  | 1 621 | 18 507 |  |  |  |
|  | SRR1513332 |  |  | 1 897 |  |  |  |  |  |  |  |
|  | SRR1513333 |  |  | 2 152 |  |  |  |  |  |  |  |
|  | SRR1513334 |  |  | 1 785 |  |  |  |  | 9 968 | 2.77 | 53.86 |
| Shark + rMATS | SRR1513329 | 206 |  | 293 |  |  |  |  |  |  |  |
|  | SRR1513330 | 256 |  | 294 |  |  |  |  |  |  |  |
|  | SRR1513331 | 217 | 5 227 | 309 |  |  | 97 | 8 539 |  |  |  |
|  | SRR1513332 | 238 |  | 306 |  |  |  |  |  |  |  |
|  | SRR1513333 | 265 |  | 331 |  |  |  |  |  |  |  |
|  | SRR1513334 | 216 |  | 284 |  |  |  |  |  |  |  |
| SUPPA2 | SRR1513329 |  |  |  |  | 333 |  |  |  |  |  |
|  | SRR1513330 |  |  |  |  | 439 |  |  |  |  |  |
|  | SRR1513331 |  |  |  | 305 | 367 | 1 010 | 3 934 |  |  |  |
|  | SRR1513332 |  |  |  |  | 532 |  |  |  |  |  |
|  | SRR1513333 |  |  |  |  | 399 |  |  |  |  |  |
|  | SRR1513334 |  |  |  |  | 549 |  |  | 1870 | 0.52 | 47.53 |
| Shark + SUPPA2 | SRR1513329 | 206 |  |  |  | 31 |  |  |  |  |  |
|  | SRR1513330 | 256 |  |  |  | 35 |  |  |  |  |  |
|  | SRR1513331 | 217 |  |  | 305 | 33 | 166 | 2 064 |  |  |  |
|  | SRR1513332 | 238 |  |  |  | 33 |  |  |  |  |  |
|  | SRR1513333 | 265 |  |  |  | 35 |  |  |  |  |  |
|  | SRR1513334 | 216 |  |  |  | 28 |  |  |  |  |  |
| SplAdder | SRR1513329 |  |  | 1 773 |  |  |  |  |  |  |  |
|  | SRR1513330 |  |  | 1 704 |  |  |  |  |  |  |  |
|  | SRR1513331 |  | 5 227 | 2 348 |  |  | 30 879 | 47 765 |  |  |  |
|  | SRR1513332 |  |  | 1 897 |  |  |  |  |  |  |  |
|  | SRR1513333 |  |  | 2 152 |  |  |  |  |  |  |  |
|  | SRR1513334 |  |  | 1 785 |  |  |  |  | 30 029 | 8.34 | 62.87 |
| Shark + SplAdder | SRR1513329 | 206 |  | 293 |  |  |  |  |  |  |  |
|  | SRR1513330 | 256 |  | 294 |  |  |  |  |  |  |  |
|  | SRR1513331 | 217 | 5 227 | 309 |  |  | 9 294 | 17 736 |  |  |  |
|  | SRR1513332 | 238 |  | 306 |  |  |  |  |  |  |  |
|  | SRR1513333 | 265 |  | 331 |  |  |  |  |  |  |  |
|  | SRR1513334 | 216 |  | 284 |  |  |  |  |  |  |  |

## References

[1] D. Belazzougui, T. Gagie, V. Mäkinen, and M. Previtali. Fully dynamic de Bruijn graphs. In S. Inenaga, K. Sadakane, and T. Sakai, editors, String Processing and Information Retrieval, pages 145–152, Cham, 2016. Springer International Publishing.

[2] A. Best, K. James, C. Dalgliesh, E. Hong, M. Kheirolahi-Kouhestani, T. Curk, Y. Xu, M. Danilenko, R. Hussain, B. Keavney, et al. Human Tra2 proteins jointly control a CHEK1 splicing switch among alternative and constitutive target exons. Nature communications, 5:4760, 2014.

[3] B. H. Bloom. Space/time trade-offs in hash coding with allowable errors. Commun. ACM, 13(7):422–426, 1970.

[4] N. L. Bray, H. Pimentel, P. Melsted, and L. Pachter. Near-optimal probabilistic RNA-seq quantification. Nature biotechnology, 34(5):525, 2016.

[5] A. Conesa, P. Madrigal, S. Tarazona, D. Gomez-Cabrero, A. Cervera, A. McPherson, M. W. Szcześniak, D. J. Gaffney, L. L. Elo, X. Zhang, and A. Mortazavi. A survey of best practices for RNA-seq data analysis. Genome biology, 17(1):13, 2016.

[6] F. Cunningham, P. Achuthan, W. Akanni, J. Allen, M. R. Amode, I. M. Armean, R. Bennett, J. Bhai, K. Billis, S. Boddu, et al. Ensembl 2019. Nucleic acids research, 47(D1):D745–D751, 2018.

[7] L. Denti, M. Previtali, G. Bernardini, A. Schönhuth, and P. Bonizzoni. Malva: genotyping by mapping-free allele detection of known variants. iScience, 2019.

[8] L. Denti, R. Rizzi, S. Beretta, G. Della Vedova, M. Previtali, and P. Bonizzoni. ASGAL: aligning RNA-Seq data to a splicing graph to detect novel alternative splicing events. BMC Bioinformatics, 19(1):444, Nov 2018.

[9] A. Dobin, C. A. Davis, F. Schlesinger, J. Drenkow, C. Zaleski, S. Jha, P. Batut, M. Chaisson, and T. R. Gingeras. STAR: ultrafast universal RNA-seq aligner. Bioinformatics, 29(1):15–21, 2013.

[10] S. Gog, T. Beller, A. Moffat, and M. Petri. From theory to practice: Plug and play with succinct data structures. In 13th International Symposium on Experimental Algorithms, (SEA 2014), pages 326–337, 2014.

[11] T. Griebel, B. Zacher, P. Ribeca, E. Raineri, V. Lacroix, R. Guigó, and M. Sammeth. Modelling and simulating generic rna-seq experiments with the flux simulator. Nucleic acids research, 40(20):10073–10083, 2012.

[12] A. Kahles, K.-V. Lehmann, N. C. Toussaint, M. Hüser, S. G. Stark, T. Sachsenberg, O. Stegle, O. Kohlbacher, C. Sander, and S. J. e. a. Caesar-Johnson. Comprehensive analysis of alternative splicing across tumors from 8,705 patients. Cancer cell, 34(2):211–224, 2018.

[13] A. Kahles, C. S. Ong, Y. Zhong, and G. Rätsch. *SplAdder*: identification, quantification and testing of alternative splicing events from rna-seq data. Bioinformatics, 32(12):1840–1847, 2016.

[14] M. Kokot, M. Długosz, and S. Deorowicz. KMC 3: counting and manipulating k-mer statistics. Bioinformatics, 33(17):2759–2761, 05 2017.

[15] G. Marçais and C. Kingsford. A fast, lock-free approach for efficient parallel counting of occurrences of k-mers. Bioinformatics, 27(6):764–770, 01 2011.

[16] R. Patro, G. Duggal, M. I. Love, R. A. Irizarry, and C. Kingsford. Salmon provides fast and bias-aware quantification of transcript expression. Nature methods, 14(4):417, 2017.

[17] R. Patro, S. M. Mount, and C. Kingsford. Sailfish enables alignment-free isoform quantification from RNA-seq reads using lightweight algorithms. Nature biotechnology, 32(5):462–464, May 2014.

[18] G. A. Sacomoto, J. Kielbassa, R. Chikhi, R. Uricaru, P. Antoniou, M.-F. Sagot, P. Peterlongo, and V. Lacroix. KISSPLICE: de-novo calling alternative splicing events from RNA-seq data. In BMC bioinformatics, volume 13, page S5. BioMed Central, 2012.

[19] S. Shen, J. W. Park, Z.-x. Lu, L. Lin, M. D. Henry, Y. N. Wu, Q. Zhou, and Y. Xing. rmats: robust and flexible detection of differential alternative splicing from replicate rna-seq data. Proceedings of the National Academy of Sciences, 111(51):E5593–E5601, 2014.

[20] A. Srivastava, H. Sarkar, N. Gupta, and R. Patro. Rapmap: a rapid, sensitive and accurate tool for mapping rna-seq reads to transcriptomes. Bioinformatics, 32(12):i192–i200, 2016.

[21] C. Sun, R. S. Harris, R. Chikhi, and P. Medvedev. Allsome sequence bloom trees. Journal of Computational Biology, 25(5):467–479, 2018.

[22] C. Sun and P. Medvedev. Toward fast and accurate snp genotyping from whole genome sequencing data for bedside diagnostics. Bioinformatics, 35(3):415–420, 2018.

[23] J. Tazi, N. Bakkour, and S. Stamm. Alternative splicing and disease. Biochimica et Biophysica Acta (BBA) – Molecular Basis of Disease, 1792(1):14 – 26, 2009.

[24] J. L. Trincado, J. C. Entizne, G. Hysenaj, B. Singh, M. Skalic, D. J. Elliott, and E. Eyras. Suppa2: fast, accurate, and uncertainty-aware differential splicing analysis across multiple conditions. Genome biology, 19(1):40, 2018.

